# Dynamic change of calcium-rich compartments during coccolithophore biomineralization

**DOI:** 10.1101/2024.11.15.623571

**Authors:** Alexander Triccas, Daniel M. Chevrier, Mariana Verezhak, Johannes Ihli, Manuel Guizar-Sicairos, Mirko Holler, Andre Scheffel, Noriaki Ozaki, Virginie Chamard, Rachel Wood, Tilman Grünewald, Fabio Nudelman

## Abstract

Coccolithophores are abundant marine phytoplankton that produce biomineralized calcite scales, called coccoliths, which sequester substantial amounts of carbon and play a significant role in biogeochemical cycles. However, mechanisms underlying the storage and transport of ions essential for calcification remain unresolved. We used ptychographic X-ray computed tomography under cryogenic conditions to visualize intracellular calcium-rich structures involved in the storage of calcium ions in the coccolithophore species *Chrysotila carterae*. During calcification, we observed a range of structures, from small electron-dense bodies within larger compartments, to denser and distributed globular compartments, before returning to small bodies once scale formation is complete. Nanobeam-scanning X-ray fluorescence measurements further revealed these electron-dense bodies are rich in P and Ca (molar ratio of ∼4:1). We infer from the dynamic nature of structures that these bodies are part of required cellular calcium ion transport pathways, a fundamental process critical for understanding the response of coccolithophores to climate change.

## Introduction

Coccolithophores are marine unicellular algae that are among the most prolific calcifiers in the oceans. They produce polycrystalline scales made of calcite called coccoliths, displaying a remarkable ability to orchestrate the multi-level assembly of nano-crystalline building blocks into a higher order structure. Coccolith production is of high importance to Earth’s biogeochemical cycle. Their formation is responsible for the sequestration of large amounts of CO_2_ within ocean sediments, forming the largest geological sink of carbon from the ocean/atmosphere reservoir and providing an important stabilizing feedback in Earth’s climate system.^1^ The continuous sinking of the scales to the ocean floor not only provides ballast for the transport of organic matter to the deep sea^2^ but is also important for the vertical gradient in sea water alkalinity^3^ and for a stabilizing feedback in Earth’s climate system.^4^ Despite the significance of coccolithophore biomineralization for our environment, we still know little about the mechanisms governing the calcification and how they can be affected by changes in ocean chemistry caused by anthropogenic activity.

Coccolith production takes place intracellularly, in a specialized compartment called the coccolith vesicle.^5^ Within, calcite crystals nucleate around the edge of an organic template called the base plate to form a ring of rhombohedral crystals called the proto-coccolith ring.^5-7^ Single crystalline units then extend sideways and outwards, mechanically interlocking as they develop. The growth and overall morphology are controlled by coccolith-associated polysaccharides,^8-10^ a constrained space for growth^11,12^ and ion gradients.^13^ Once complete, coccoliths, with size in the 1-10 micrometer range, are exocytosed and, together with previously formed scales, form an exoskeleton around the cell called the coccosphere.^14^ The cellular mechanisms that control different aspects of coccolith mineralization are still left to be clarified. One particular area that is not well understood is the mass transport of ions across the cell to the mineralization site. A significant quantity of calcium ions are required to produce condensed mineral phases, which raises questions on how coccolithophores, and eukaryotic cells in general, avoid cytotoxicity. Calcium concentration in the cytosol must be kept around 100 nM to prevent toxicity and eventual cell death^15^. For calcite to precipitate, concentrations within the coccolith vesicle must be raised to 100-200 µM.^16^ Cellular pathways are therefore required to transport calcium ions to satisfy the demands of calcification, without raising cytosolic concentrations to fatal levels.

Several biomineralizing organisms, including corals, foraminifera and sea urchin larvae, source calcium from the surrounding seawater by endocytosis.^17-19^ Once inside the cell, a hydrated amorphous calcium carbonate phase forms within intracellular vesicles, which are transported to the mineralization site.^20^ In the case of coccolithophores, calcium enters the cell via ion channels on the plasma membrane, before being loaded into endomembrane compartments.^21,22^ For the coccolithophore *Chrysotila carterae*, calcium ions are delivered to the coccolith vesicle in the form of coccolithosomes, which are electron-dense ∼25 nm calcium-associated polysaccharide complexes that originate in the Golgi cisternae.^5,23^ Once inside the vesicle, they provide calcium ions for nucleating and growing calcite crystals.^6,24^

It is unclear how calcium ions are transported across the cell into the Golgi network in order for coccolithosomes to form. Large intracellular pools of calcium and polyphosphates resembling acidocalcisomes, which are storage organelles within numerous prokaryotic and eukaryotic cells,^25^ were identified in coccolithophores.^26,27^ The role of these structures in calcification is still debated. While they have been initially proposed to be part of a calcium storage and transport pathway involved in coccolith formation,^26,27^ subsequent fluorescent studies suggest that the composition, their distribution, and calcium content did not change during coccolith production.^28^ These findings led to the hypothesis that such compartments may function more passively as storage organelles to control and equilibrate calcium concentrations inside the cell.^28^ However, calcium-rich structures have mostly been characterised from single snapshots taken across cryogenically or chemically-fixed coccolithophore cells or with confocal microscopy techniques that have limited spatial resolution.^26-28^ As a result, any dynamic changes that may occur while cells are calcifying could remain undetected.

To investigate the potentially hidden dynamic pathways that transport calcium through the cell during calcification, we determined how calcium-rich structures change in distribution and density during this process. For this, whole cells of the coccolithophore *C. carterae* were imaged using ptychographic X-ray computed tomography in cryogenic conditions (cryoPXCT). X-ray ptychography is a quantitative 3D microscopy approach, which provides access to the mass density at the sub-micrometer scale.^29-32^ Additionally, nanobeam-scanning X-ray fluorescence (nXRF) was used to determine the calcium content of these dense structures and assess their overall elemental composition. We identified that the distribution of calcium ions in intracellular structures changed across the window in which calcification was taking place. These findings will strengthen our overall understanding of the calcium transport pathways employed by coccolithophore cells to facilitate the demands of coccolith production.

## Results

### Calcium-rich structures in *C. carterae* cells

Actively calcifying cells of the coccolithophore *C. carterae* were treated with 0.1 M EDTA for 5 min in order to remove their external coccoliths, before being incubated in low calcium medium for 24 h to inhibit mineralization. Calcium was then reintroduced to the medium to restart calcification. After 4, 8, 12 and 24 h of incubation, cells were loaded into glass capillaries, plunge-frozen in liquid ethane, and analysed using cryoPXCT at -180 °C. This synchrotron-source based technique provided 3D reconstructions with quantitative electron density information and spatial resolution in the 10-100 nm range.^29-31^ Data segmentation was achieved based on the electron density of the cellular structures, evidencing extracellular and intracellular coccoliths, along with the intracellular organelles and compartments. The longer cells were incubated in calcium-replete medium, more coccoliths were produced and the coccosphere became more developed. A degree of variability in the rate of formation of the coccosphere was observed at each incubation time, and therefore the cells were categorized according to the stages of formation of their coccospheres, going from stage 1 (only few coccoliths formed) to stage 4 (most of the coccosphere complete). This is represented in Figure. 1, where an example of one cell at each state of formation is shown. In stage 1, few external coccoliths were present (in yellow, Figure 1a-b and Supplementary Movie 1). In stage 2, 15-20 external coccoliths were spread around the cell surface (in yellow, Figure 1c-d and Supplementary Movie 2). In stage 3, the coccosphere was more complete, with adjacent coccoliths starting to connect (in yellow, Figure 1e-f and Supplementary Movie 3). At these three stages of coccosphere formation, the majority of cells contained intracellular coccoliths (coloured as red for sake of clarity in Figure 1); indicating calcification was actively taking place within. Finally, in stage 4 large parts of the coccosphere were complete (Figure 1g-h, yellow). While a forming coccolith can be observed in the cell shown in Figure 1g-h (see also Supplementary Movie 4), three cells out of six at this stage contained no intracellular mineral structures (Supplementary Movie 5 depicts the reconstructions of all 6 cells in this stage). In all cases, additional features were visible alongside the intracellular and external coccoliths, including structures with electron density between 0.42 and 0.62 n_e_Å^-3^ (dark blue, Figure 1) and organelles and compartments with electron density between 0.37-0.40 n_e_Å^-3^ that were either empty (Figure 1, turquoise) or containing the denser structures with electron density between 0.42 and 0.62 n_e_Å^-3^ (Figure 1, pink).

**Figure 1.**
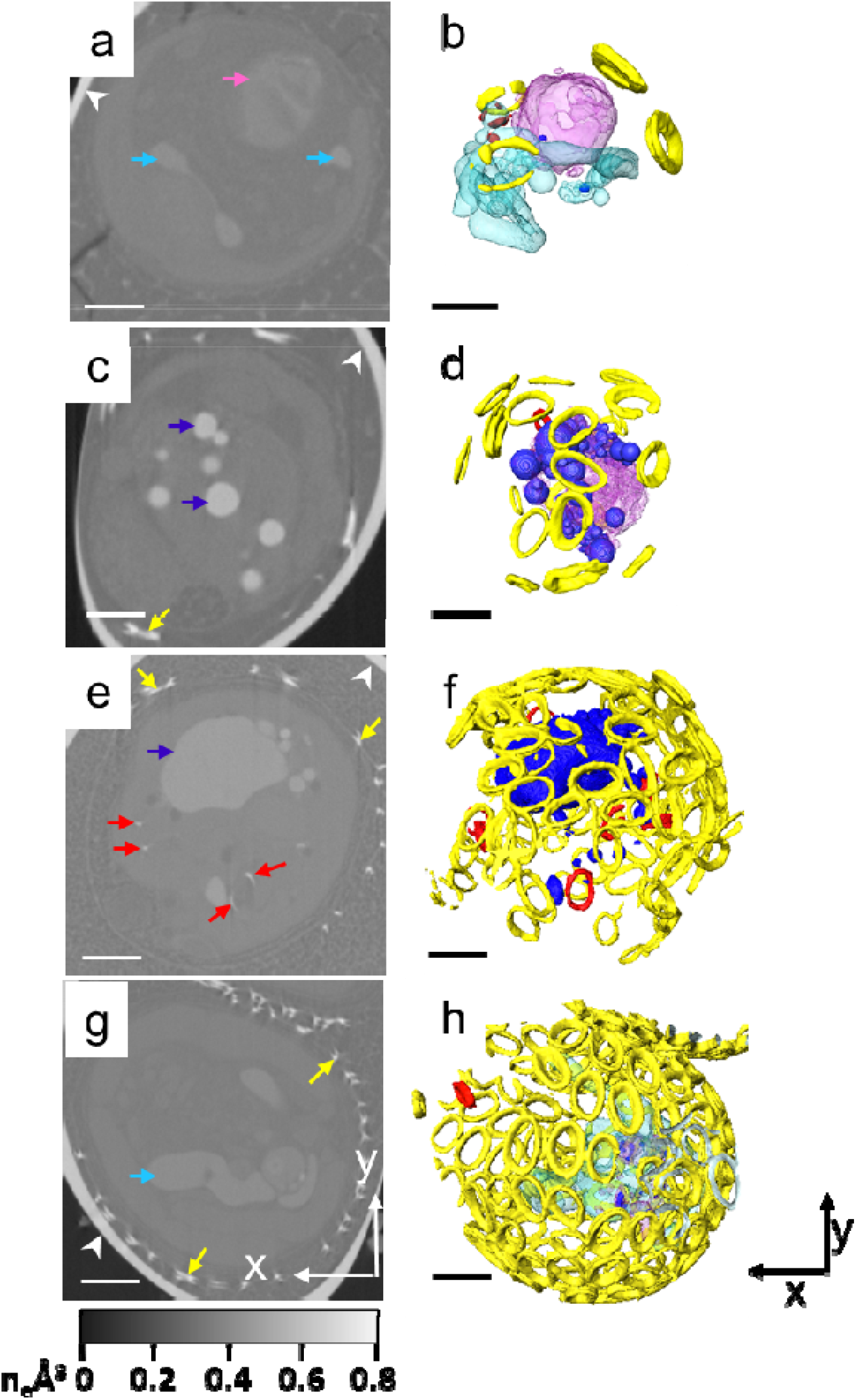
CryoPXCT tomograms of calcifying *C. carterae* cells frozen at different stages of calcification. Left column: 2D slice through the volume of individual cells. Right column: respective isosurface-rendered visualisation. Visible are: extracellular coccoliths (yellow arrows in c-g and colored in yellow in b, d, f and h); intracellular coccoliths (red arrows in e and red in d,f, and h); organelles with electron density between 0.37 and 0.42 n_e_Å^-3^ (turquoise arrows in a and g and colored in turquoise in b; pink arrows in a and colored pink in b and d); and structures with electron density above 0.42 n_e_Å^-3^ (dark blue arrows in c and e and colored dark blue in a, b, d, and f). (a,b) Stage 1. (c, d) Stage 2. (e, f) Stage 3. (g, h) Stage 4. Arrowheads indicate the glass capillary used to store the sample. Scale bars = 2 µm.

### Composition of the dense intracellular deposits in *C. carterae*

As the electron density for most cellular bodies is reported^33^ to be up to 0.35 n_e_Å^-3^, we hypothesize that that the structures with values of 0.37 n_e_Å^-3^ and higher contain heavier elements, with calcium being a likely candidate considering that it is stored inside intracellular compartments.^26,27^ To verify this hypothesis, we determined the elemental composition of the dense intracellular structures by nano-scanning X-ray fluorescence (nXRF). Subsequently, using our cryoPXCT data, we analysed their distribution inside the cells across the different stages of calcification to determine if any dynamic changes in the intracellular localization of calcium during coccosphere formation could be observed.

*C. carterae* cells were collected after 4 h in Ca-replete medium, dried on substrate and then imaged by nano-scanning X-ray fluorescence (nXRF). In this approach, the signal is summed along the beam propagation direction, so that the presented image corresponds to an integration of the signal along the cell thickness. Differential phase contrast (DPC) was collected simultaneously with nXRF to provide a reference image of the coccolithophore cell measured (See Materials and Methods). DPC maps of two cells are presented in the first column of Figure 2a-b both showing one associated coccolith. In the two subsequent columns are elemental maps of Ca and P (Supplementary Fig. 1 shows representative fitting examples for nXRF datasets), which have XRF signals pertaining to coccolithophore cells (Supplementary Fig. 2 shows S Kα XRF signals pertaining to concentrated cellular material observed with DPC). The Ca Kα XRF signal is strongly associated with coccolith materials (Figure 2) but can also be observed in distinct regions of the cell (Supplementary Fig. 2). These regions are also dominated by the P Kα XRF signal (Figure 2b and Supplementary Fig. 2) and are comparable in appearance to the roundish structures in cryoPXCT tomograms (Figure 1c), though slightly larger due to flattening effects from drying. Cell 2 (Figure 2b) contains many more dense compartments than Cell 1; considering that cells at Stage 2 are characterized by larger numbers of dense bodies that Stage 1 cells (Figure 1), it suggests that Cell 2 was at a later stage of calcification (e.g., see Stage 2 cryoPXCT in Figure 4). It is expected that lower electron density features (e.g., 0.37-0.40 n_e_Å^-3^) are less discernible with nXRF and that their structural preservation is likely lost.

**Figure 2.**
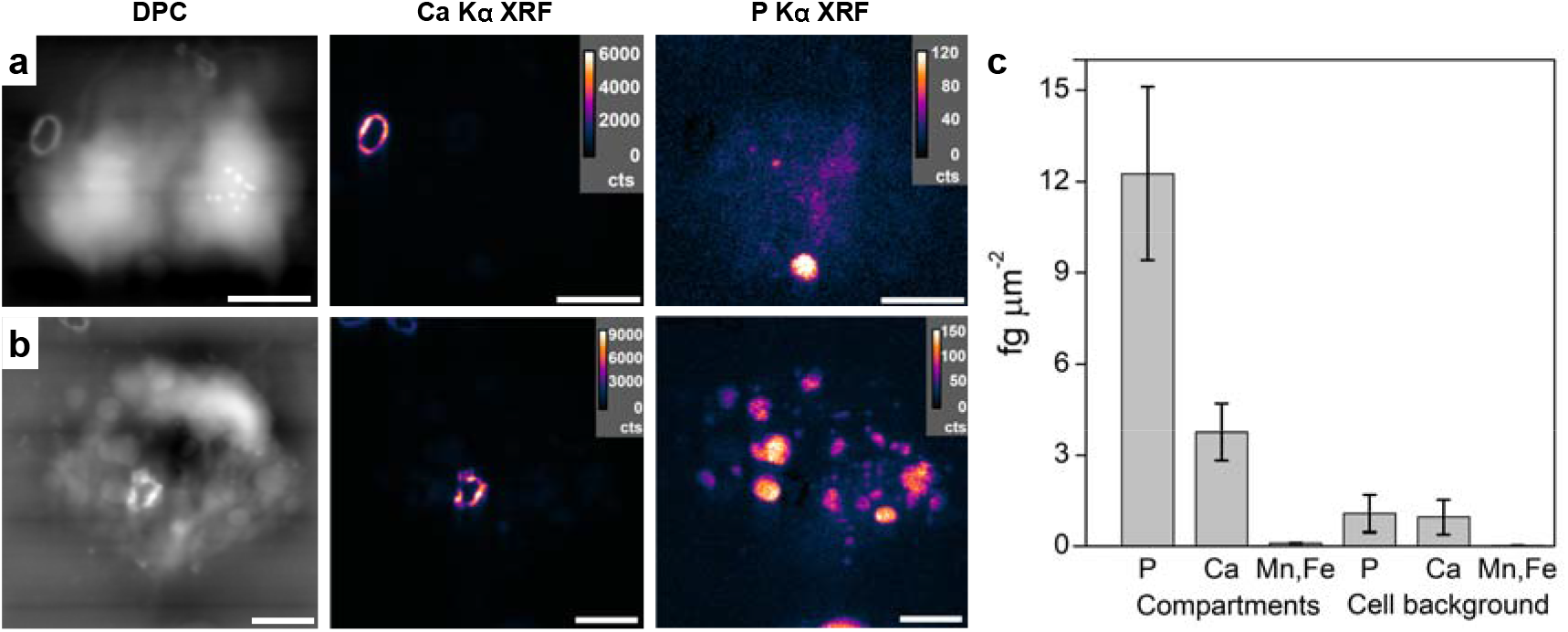
Nano-scanning X-ray fluorescence measurements of dried *C. carterae cells*. Differential phase contrast (DPC) and X-ray fluorescence maps (XRF) maps of *C. carterae* (a) Cell 1 and (b) Cell 2 harvested after 4 h in Ca-replete medium and dried on substrate. Principal elements of interest from XRF data are shown with linear-based intensity from detector counts (cts). (c) Semi-quantitative elemental quantities of selected elements based on XRF intensities. Scale bars: 4 µm.

**Figure 3:**
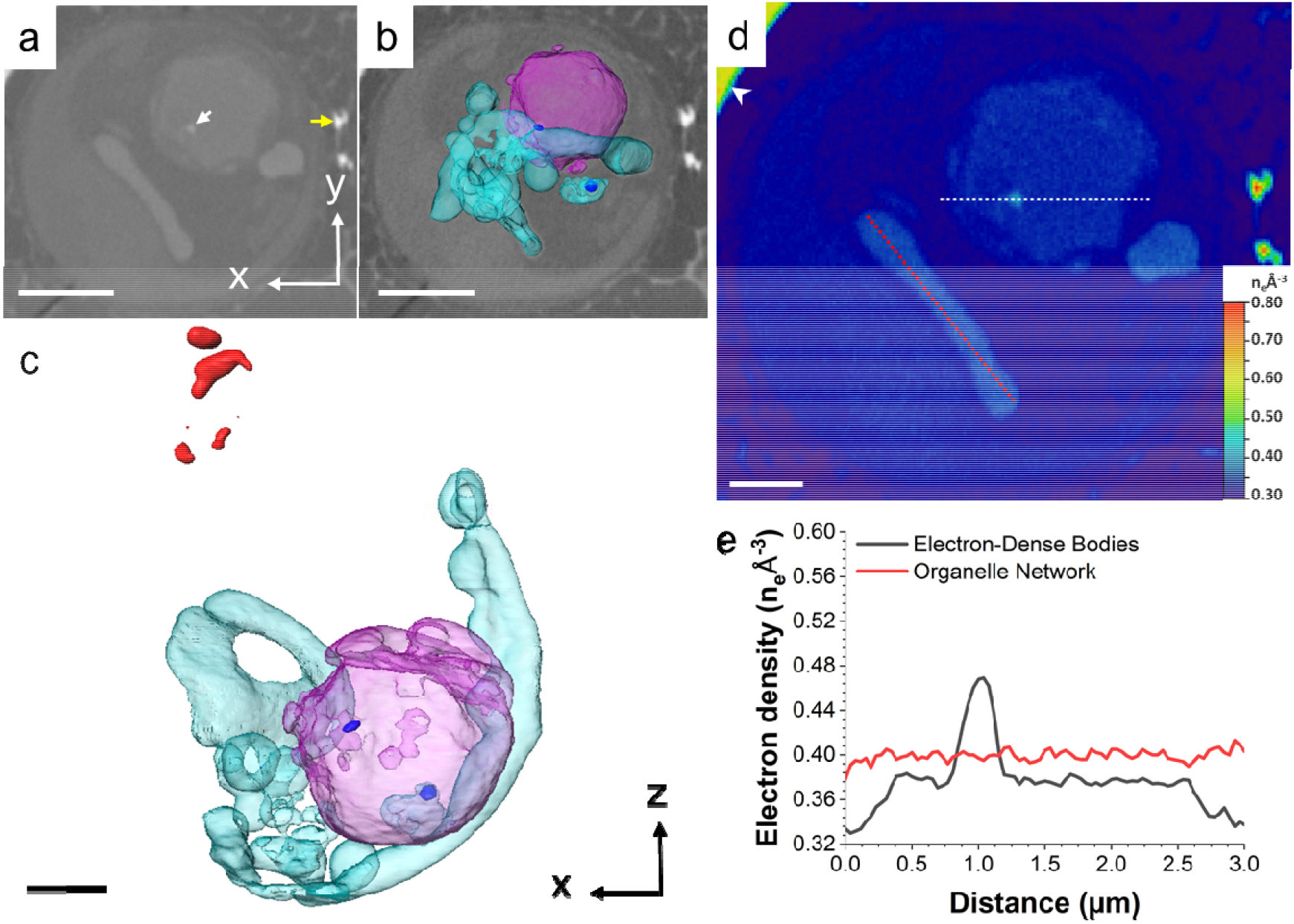
CryoPXCT tomogram of intracellular structures within a *C. carterae* cell at stage 1. (a) 2D slice through the volume of the cell. White arrow: electron dense body. Yellow arrow: mature coccolith on the surface of the cell. The 3D isosurface-rendered visualization of the intracellular electron-dense structures is shown overlaid on the 2D slice (b), and in more detail in (c). Visible are: a spherical compartment (pink) with electron density ∼0.37 n_e_Å^-3^ containing bodies with electron density above 0.43 n_e_Å^-3^ (dark blue); an organellar network with electron density of 0.40 n_e_Å^-3^ (turquoise); and potential calcite deposits of forming coccoliths with an electron density above 0.70 n_e_Å^-3^ (red). (d) Electron density map of (a). Arrowhead indicates the wall of the glass capillary used to store the sample. (e) Electron density line profiles of the areas marked by the white and red lines in (d). Scale bars: 2 µm (a,b), 1 µm (c,d).

**Figure 4:**
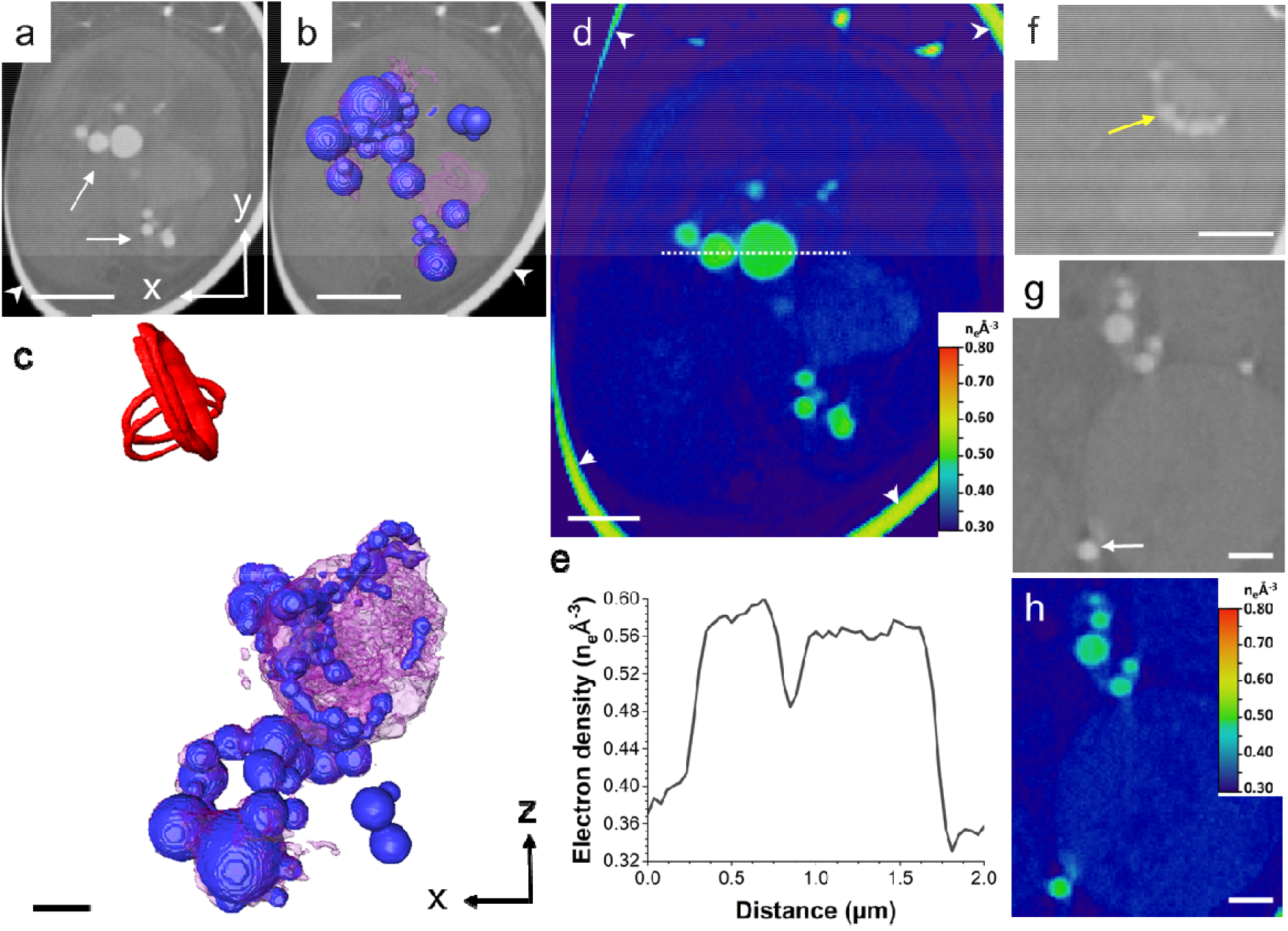
CryoPXCT tomogram of intracellular structures within a *C. carterae* cell at stage 2. (a) 2D slice through the volume of the cell. The 3D isosurface-rendered visualisation of all electron-dense structures is shown overlaid on the 2D slice (b) and in more detail in (c). Visible are: Dense bodies with electron density between 0.56-0.60 n_e_Å^-3^ distributed across the cell (arrow in a, dark blue b,c), intracellular structures with electron density 0.37 n_e_Å^-3^ (pink) and forming intracellular coccoliths (red). (d) Electron density map of (a). (e) Electron density line profile of the area marked by the white line in (d). (f-g) Two 2D slices from different areas of the cell showing in higher detail electron-dense bodies (yellow arrow in (f) and white arrow in (g)) inside the compartment depicted in pink in (c). (h) Electron density map of (g). Scale bars: 2 µm (a,b), 1 µm (c-h). Arrowheads indicate the wall of the glass capillary used to prepare the sample.

The semi-quantitative elemental composition (standard reference calibrated, substrate background- and area-corrected signals) of the denser features and the cell background is presented in Supplementary Table 1 and summarized in Figure 2c. Most notable is the ∼11-fold increase of P and ∼4-fold increase of Ca in dense compartments compared to the cell background. Other elements, such as Mn and Fe (Supplementary Fig. 3), are lower in concentration compared to Ca and P and likely have little contribution to the electron density values measured from cryoPXCT data. In addition to the increase of P and Ca in the compartments, the relative ratio of these quantities changes from a P/Ca of ∼1.1 in the cell background to ∼3.3, converting mass to eq. moles, mol P / mol Ca of ∼4.2. Altogether, this analysis shows that dense compartments in calcifying *C. carterae* cells contain a high concentration of P in addition to Ca.

### Stage 1 – Start of coccosphere formation

Following the identification of electron-dense structures as Ca-P-rich bodies by nXRF, we analysed in detail their dynamics as a function of development stage of the coccosphere using cryoPXCT. One cell at stage 1 is shown in Figure 3 and Supplementary Movie 1. A 2D cross-section across the cell (Figure 3a) and 3D visualisation (Figure 3b and c) highlight the presence of an organellar network (turquoise) with a uniform electron density of 0.40 n_e_Å^-3^, suggesting that Ca^2+^ ions were homogeneously distributed inside (Figure 3d and e). We postulate that this organellar network corresponds to the endoplasmic reticulum, consistent with its morphology and function in calcium storage.^34^ Juxtaposed to this organellar network was one spherical compartment ∼3 µm in diameter, with an electron density of 0.38 n_e_Å^-3^. This compartment contained two ∼350 nm bodies where electron density reached 0.47 n_e_Å^-3^, which, together with the nXRF data, indicates they are Ca-P-rich bodies (Figure 3a, white arrow; dark blue in Figure 3b-c; electron density shown in 3d-e). Structures with electron density of ∼0.70 to 0.75 n_e_Å^-3^ (Figure 3c in red and Supplementary Fig. 4) were also present. These values are more consistent with that of a mineral phase,^35^ indicating that it is composed of calcium carbonate (Supplementary Fig. 4). It is conceivable that these represent the onset of calcite deposition during the formation of the protococcolith, or a malformed coccolith. A second cell at stage 1, with only one external and no intracellular coccoliths, displayed similar distribution of calcium-rich structures, only with a larger number of dense bodies of electron density of 0.48 n_e_Å^-3^ contained within compartments of electron density of 0.37 n_e_Å^-3^ (Supplementary Fig. 5 and Supplementary Movie 6).

### Stage 2 – Early stage of coccosphere formation

Two cells were analyzed at stage 2 displaying multiple external coccoliths, as well as intracellular ones being formed. The first cell is depicted in Figure 1c and d, Figure 4 and Supplementary Movie 2, and the second cell is shown in Supplementary Fig. 6 and Supplementary Movie 7. The morphology and distribution of the electron-dense Ca-P-rich bodies were vastly different compared to those present at stage 1: they were significantly more abundant, had globular shapes, and were clustered into a network-like structure (Figure 4a, arrow, and 4b and c in blue), resembling those described by cryo-X-ray tomography.^26^ The size of the bodies varied, with some as small as 350 nm, while others reached up to 1.7 µm. Their electron densities had also significantly risen compared to stage 1, with maximum values now between 0.57 and 0.61 n_e_Å^-3^ (Figure. 4d and e). When bodies were localized inside lower density structures, they were always close to the periphery (Figure 4f, yellow arrow), in some cases seemingly budding off (Figure 4g, white arrow). Electron density values in these bodies were within the same range as those freely distributed (Figure 4h). It is conceivable that they could have been produced within the larger organelles, before budding off. While the second cell displayed similar globular structures, they were lower in number when compared to the first cell at this stage (Supplementary Fig. 6 and Supplementary Movie 7). The organellar network with uniform electron density, namely the endoplasmic reticulum, was not distinguishable at this stage. A likely cause is the depletion of its Ca^2+^ content, decreasing the electron density and making it undistinguishable from the neighbouring intracellular structures and organelles with electron density up to 0.35 n_e_Å^-3^.

### Stage 3 – Mid stage of coccosphere formation

Two cells with approximately half of the coccosphere complete were classified as in stage 3 (Figure 1e and f and Figure 5, yellow arrows in Figure 5a; Supplementary Movie 3; additional cell at this stage in Supplementary Fig. 7 and Supplementary Movie 8). Intracellular coccoliths at various stages of maturity were present (red arrows in Figure 5a and segmented in red in Figs. 5c and d; Supplementary Fig. 7b and c). A significant portion of the cell was dominated by one large electron-dense body, around 2-3 µm in diameter at its thickest (Figure 5a and c-d and Supplementary Fig. 7). Electron density was consistent across the body, around 0.44 n_e_Å^-3^ (Figure 5e and g and Supplementary Fig. 7d and e). Additionally, several smaller electron-dense bodies ranging in size between 160-900 nm were present, seemingly extending away from the larger body in an extended network (Figure 5d and f and Supplementary Fig. 7a-c). Their electron density varied, with values either being 0.44 n_e_Å^-3^, the same as the larger body, or slightly higher, around 0.52 n_e_Å^-3^ (Figure 5e-h). Compared to those in stage 2 (Figure 4), the electron-dense bodies in the cells at stage 3 had lower electron density values and less defined morphologies. Given the morphological similarity of the large body to the organelle network present at stage 1, we suggest that its appearance signifies the reformation of the disperse calcium stores, with the smaller dense bodies coalescing to form the larger structure.

**Figure 5:**
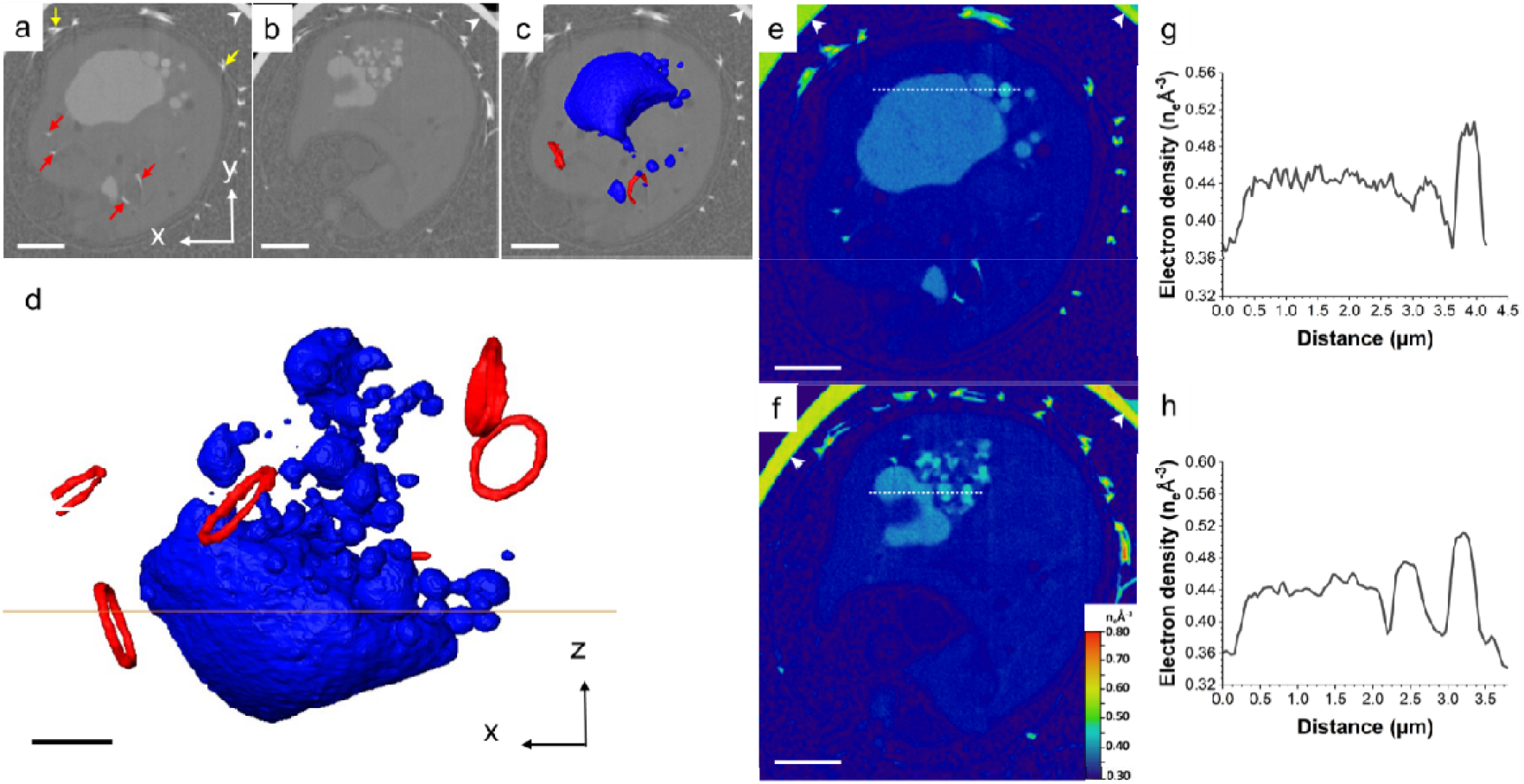
CryoPXCT tomogram of intracellular structures within a *C. carterae* cell at stage 3. (a) and (b) 2D slices through the volume of the cell. Red arrows: forming coccoliths. Yellow arrows: Mature coccoliths on the surface of the cell. The 3D isosurface-rendered visualisation of the intracellular electron-dense structures overlaid on the 2D slice from (a) is shown in (c) and in more detail in (d). Visible are: Dense bodies with electron density between 0.44-0.52 n_e_Å^-3^ (dark blue) and nascent intracellular coccoliths (red). (d) and (f) electron density maps of (a) and (b), respectively. (g) Electron density line profile of the area marked by the white line in (e). (h) Electron density line profile of the area marked by the white line in (f) Scale bars: 2 µm (a-c; e-f), 1 µm (c). Arrowheads indicate the wall of the glass capillary used to store the sample.

### Stage 4 – Final stage of coccosphere formation

Six cells were analyzed at stage 4, where large sections of the coccosphere are completed. Intracellular coccoliths were only present in three out of these cells, indicating that calcification may have been slowing down at this time point, resulting in some cells not producing coccoliths.^36^ One representative cell where no intracellular coccoliths were present, along with the corresponding 3D visualization, is shown in Figure 6 and Supplementary Movie 9.

**Figure 6:**
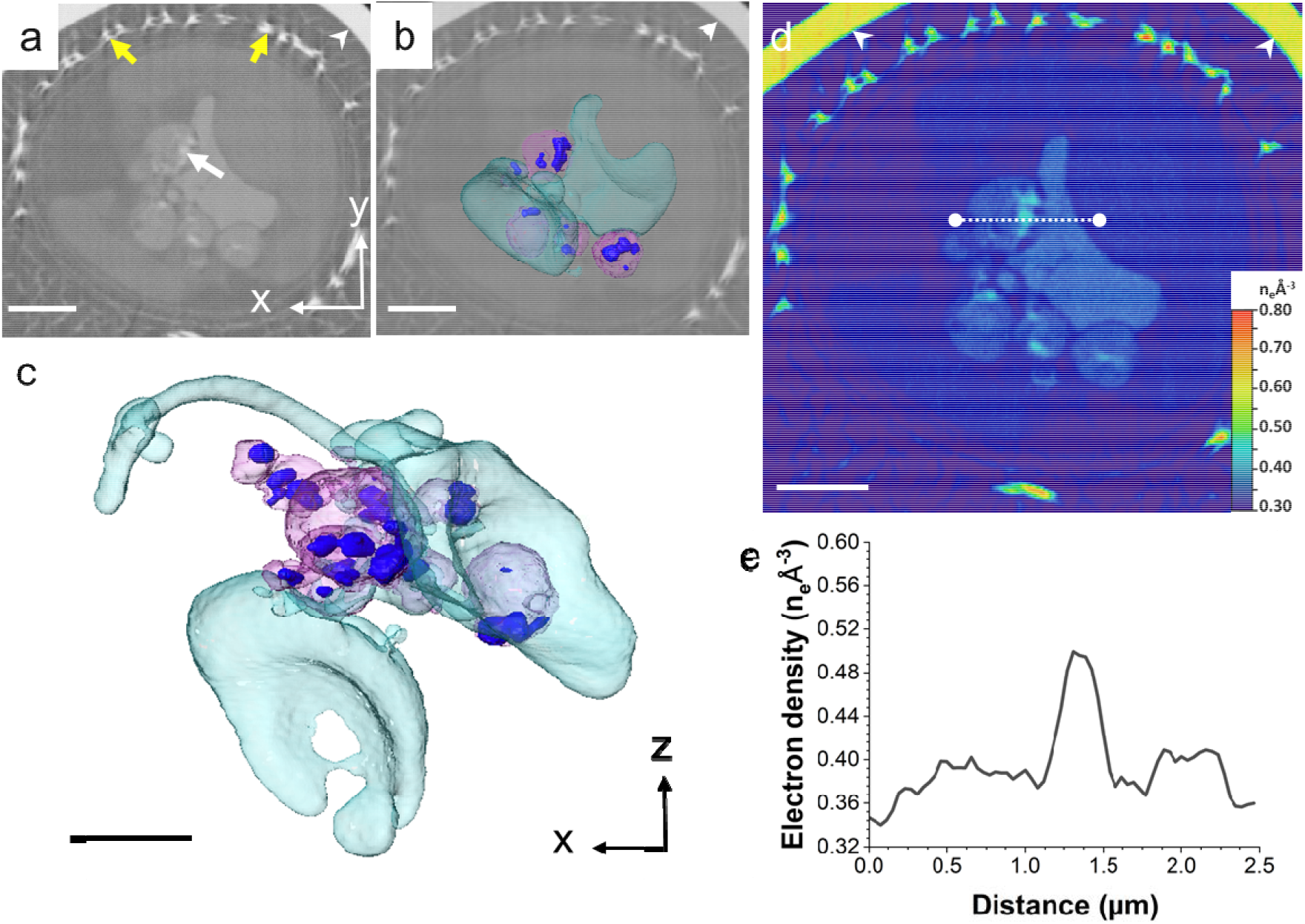
CryoPXCT tomogram of intracellular structures within a *C. carterae* cell after 24 h in calcium-replete medium. (a) 2D slice through the volume of the cell. White arrows: dense bodies with electron density around 0.50 n_e_Å^-3^. Yellow arrows: extracellular mature coccoliths. The 3D isosurface-rendered visualization of the intracellular electron-dense structures is shown overlaid on the 2D slice (b), and in more detail in (c). Visible are: A spherical compartment (pink) with electron density ∼0.37 n_e_Å^-3^ containing bodies with electron density above 0.43 n_e_Å^-3^ (dark blue); an organelle network with electron density of 0.40 n_e_Å^-3^ (turquoise). (d) Electron density map of (a). (e) Electron density line profiles of the areas marked by the white line in (d). Scale bars: 2 µm (a-c), 1 µm (d). Arrowheads indicate the wall of the glass capillary used to store the sample.

Extracellular mature coccoliths are visible (Figure 6a, yellow arrows) and several 250-500 nm-sized Ca-P-rich bodies with electron densities of 0.49-0.51 n_e_Å^-3^ were present (white arrow in Figure 6a and dark blue in Figure 6b and c; electron density displayed in Figure 6d and e), notably bound within larger compartments with electron densities of 0.37-0.38 n_e_Å^-3^ (pink, Figure 6b and c and Figure 6d and e). Additionally, the endoplasmic reticulum, displaying consistent electron density values of 0.40 n_e_Å^-3^, could be observed (Figure 6b and c, in turquoise, and Figure 6d-e). Overall, the size, morphology and distribution of structures with electron density above 0.37 n_e_Å^-3^ was similar to that observed in the cells in stage 1 (Figure 3). A similar distribution of intracellular structures and Ca-P-rich bodies was present in other cells imaged at the same time point, both with and without intracellular coccoliths (Supplementary Fig. 8 and Supplementary Movies 4 and 5).

From the electron density distribution in the intracellular compartments across the different stages of formation of the coccosphere, we infer that calcium ions were distributed in intracellular structures in two different forms across the time window in which cells were actively calcifying. At the start (stage 1) and towards the end of the formation of the coccosphere (stage 4), calcium ions were dispersed in an organellar network (turquoise in Figs. 3 and 6). We posit that this network corresponds to the endoplasmic reticulum. In addition, they were being concentrated in small bodies contained within larger compartments that were in proximity with the endoplasmic reticulum network. Between these time points, calcium storage switched to being heavily concentrated exclusively inside globular, electron-dense bodies. This transition indicates that ion concentrations can be dynamically altered in the calcium-rich structures to fulfil the demands of coccosphere formation.

## Discussion

The changes in the morphology, intracellular distribution and electron density of the calcium-rich bodies and compartments during coccosphere formation reported in this study demonstrates the dynamic nature of these structures in the coccolithophore *C. carterae*. At the start of the formation of the coccosphere – when the first coccoliths are being produced – calcium ions were homogeneously dispersed in the endoplasmic reticulum and concentrated in ∼300 nm sized Ca-P-rich bodies located within spherical compartments. As the formation of the coccosphere proceeded, the Ca-P-rich bodies increased in size, electron density and number, resembling those reported in *C. carterae* cells in previous studies.^26,28^ The endoplasmic reticulum was no longer distinguishable, indicating it had electron density values similar to those of the adjacent cellular structures, likely because of decreased calcium ion content. As the coccosphere neared completion, the distribution of dense bodies resembled the state of the earliest stage.

For the cryoPXCT measurements, individual capillaries were prepared containing cells that were harvested after spending different periods in calcium replete medium: 4h, 8h, 12h and 24h. Notably, the capillaries frozen after 8h and 12h contained cells at different stages of formation of the coccosphere: one cell at stage 1 (Supplementary Fig. 5) and one at stage 2 (Figure 4) were found after 8h; one cell at stage 2 (Supplementary Fig. 6) and two cells at stage 3 (Figure 5 and Supplementary Fig. 7) were present after 12h. These observations show that each cell likely started calcifying at different rates after being transferred from calcium-deplete to calcium-replete medium. Importantly, the distribution of calcium ions between the endoplasmic reticulum and Ca-P-dense bodies was related to the activity of the formation of the coccosphere, rather than the time spent in calcium-replete medium. Therefore, we attribute the distribution of calcium in these structures, as well as their sizes and morphologies, to not be primarily influenced by external ion concentrations. Instead, we suggest calcium distribution across the endoplasmic reticulum and Ca-P-rich bodies correlates to the calcification process, namely the mobilization and transport of calcium ions to the coccolith vesicle for coccolith formation. Calcium ions enter the cell via voltage-gated channels,^21,22^ where they are proposed to be partitioned directly into endomembrane-derived organelles, among them the endoplasmic reticulum, to avoid cytosolic concentrations interfering with calcium signalling.^34^ The association of the endoplasmic reticulum, rich in calcium ions, with the 1-2 µm spherical compartments containing the dense Ca-P-rich bodies suggests a connection between these two organelles. It is conceivable that calcium ions are transported from the endoplasmic reticulum to the spherical compartments where they form complexes with polyphosphates^27,37^ to form dense bodies with a P to Ca molar ratio of ∼4:1, according to our nXRF measurements, enabling high concentration storage of calcium ions. The dynamics between these two organelles changes as a function of coccosphere formation, likely in response to the high demand for calcium ions for coccolith formation. In this case, the Ca-P-rich bodies increase in size and number, consuming calcium ions stored in the endoplasmic reticulum, from where they are further mobilized for coccolith formation.^37^ As the coccosphere nears completion, coccolith production slows down, decreasing the demand for calcium ions which accumulate again in the endoplasmic reticulum. This model is summarized in Figure 7.

**Figure 7.**
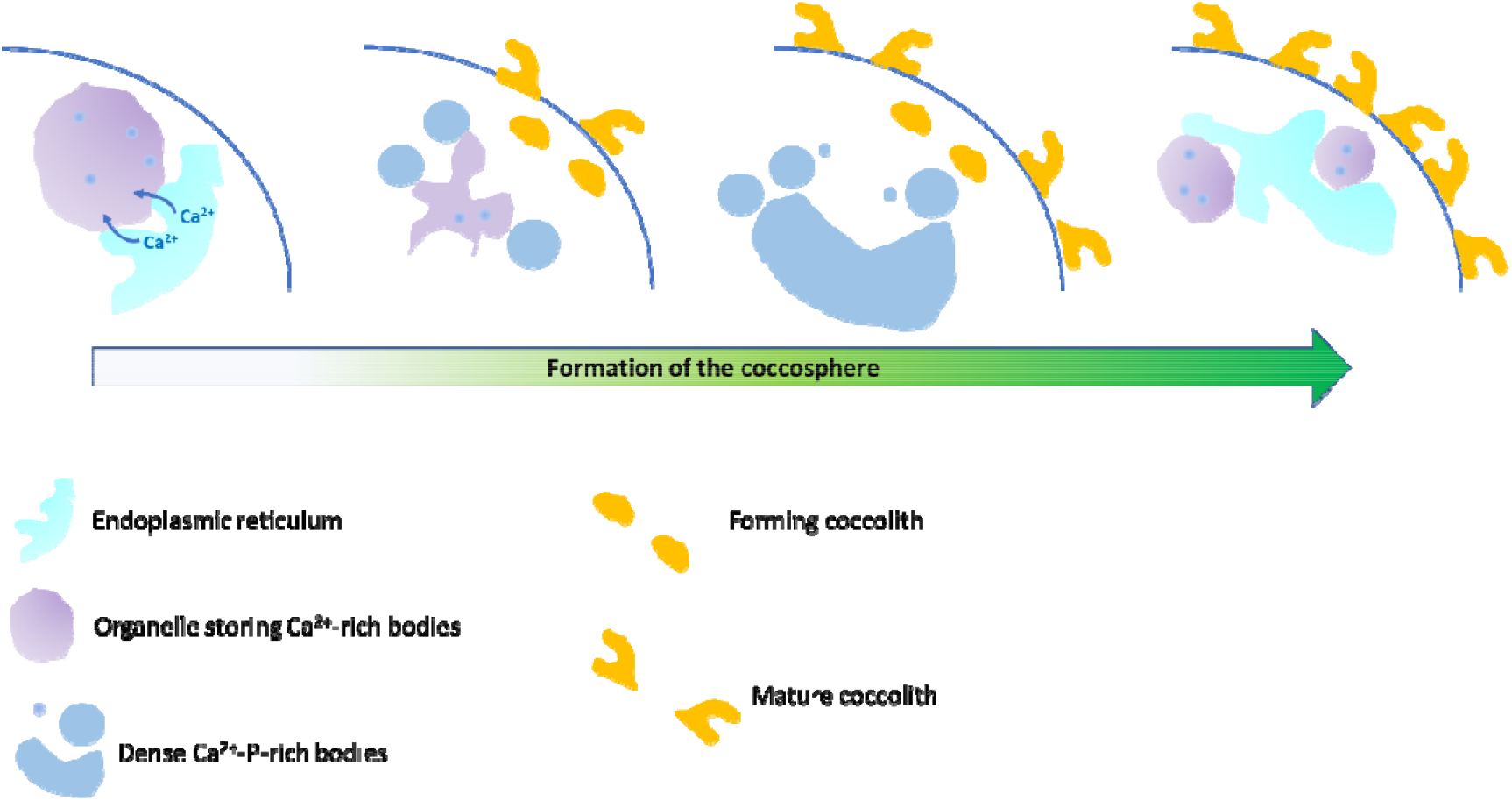
Dynamics of calcium-rich compartments during coccosphere formation in *C. carterae*. Schematic of the proposed changes in the calcium storage compartments of coccolithophores during the formation of the coccosphere.

It has been suggested that calcium-rich compartments and Ca-P-rich dense bodies are present in coccolithophores independent of calcification activity, possibly providing an alternate function to the cell.^28^ Spectroscopic studies, however, indicate that ions from the dense bodies become incorporated into the coccolith crystal lattice, which requires a mechanism to transport calcium from the Ca-P-rich bodies into the coccolith vesicle. Our study shows that the dense bodies fluctuate in electron density and number across coccosphere development, supporting the hypothesis that they alter their ion capacity and function as dynamic calcium reservoirs in response to the demands of biomineral formation.^27^This dynamic function could facilitate the transfer of calcium from the dense bodies to the coccolith vesicle. This pathway requires the transport of ions through the Golgi cisternae, where coccolithosomes – 25 nm calcium-polysaccharide complexes that deliver calcium ions to the coccolith vesicle – originate.^23,24^ The presence of these particles could not be confirmed by cryoPXCT as they are below the resolution limit of this imaging technique.

Due to their resemblance to acidocalcisomes, it appears that calcium-rich pools existed prior to coccolithophore calcification and may have been adapted to store and transport the large quantities of calcium to facilitate mineral formation. Intracellular calcium pools are not a common feature of the biomineralization process, absent in most mineralizing organisms, with ions instead transported alongside carbonate in ACC-containing vesicles derived from the endocytosis of seawater.^18,34^ Calcium-rich pools have been reported in dinoflagellates, which, like coccolithophores, transport calcium ions into the cell via gated channels.^38^ In dinoflagellates, the calcium-rich bodies are generally associated with extending the mineral front, similar to the ACC-vesicles in other biomineralizing organisms.^18,20,38^ It is possible that alternate transport pathways may have developed in coccolithophores as a result of crystallization occurring in an intracellular environment. It is also possible that several undetected pathways for ion delivery might exist, varying from species to species, with possible influence from the external cellular environmental conditions.

In conclusion, the findings reported in this study reveal new insights into the role of intracellular calcium-rich structures in coccolithophore calcification. By characterising the calcium distributions during *C. carterae* calcification, we have determined that calcium-storage compartments and Ca-P-rich bodies act as dynamic reservoirs which are able to fluctuate the quantity of ions contained within. These observations indicate that such structures are integral components of the transport pathways responsible for shuttling calcium ions across the cell during calcification. They therefore play an important role in the storage and mobilization of these ions without posing a risk of cytotoxicity.

## Experimental

### Culturing

Cultures of *Chrysotila carterae* (944/6) were obtained from the Culture Collection of Algae and Protozoa (CCAP), Oban, Scotland. Cultures were incubated at 18 °C on a 12 h:12 h light-dark cycle. Cells were grown in natural seawater from St Abbs Marine Station supplemented with Guillard’s (F/2) Marine Water Enrichment Solution (Sigma Aldrich) and with a penicillin/streptomycin/neomycin antibiotics solution (Fisher BioReagents). Cultures were maintained by subculturing 1 mL of culture into 30 mL fresh medium every 2 weeks. Cell growth was measured using a Leica optical microscope and Neubauer haemocytometer.

### Cryo-ptychographic nanotomography (cryo-PXCT)

#### Preparation of *C. carterae* cells

*C. carterae* cells were prepared by decalcifying 5 mL of culture in its exponential phase (5 days after subculturing) with 0.1 M pH 8.0 EDTA to remove external coccoliths. Cells were washed twice in culture medium to remove residual EDTA, then left overnight in low-calcium (100 µM) Aquil artificial seawater to prevent further coccolith growth and recalibrate calcification. A 0.1 M CaCl_2_ x 2H_2_O (100 µL) solution was then added to raise [Ca^2+^] back to seawater levels (10 mM) and left for 24 h to provide sufficient time for enough coccoliths to form to fill the entire coccosphere. Cells were in 1 mL aliquots after 4, 8, 12, and 24 h in calcium-replete medium, centrifuged at 1,500 g, and concentrated to a ∼50 µL volume.

#### Plunge-freezing

Capillaries with tips 10 µm wide were produced using a Flaming-Brown capillary puller (P-2000, Sutter Instruments) and mounted in OMNY pins^39^ with UV resin. The concentrated solution of cells at each calcifying timepoint were loaded into the capillary tips from the accessible backside. The cells were then further sedimented into the very tip of the capillary by a small settling period or by mild centrifugation (1,120 g for 240 s). The capillaries were rapidly plunge-frozen in liquid ethane using a vitrification robot (FEI Vitrobot Mark VI). The sample chamber was maintained at 21 °C and 100% humidity. Particular care was taken to minimize the time lag between sample preparation and sample freezing. The samples were stored at cryogenic temperature under liquid nitrogen and transferred to the beamline using a transport Dewar to ensure a continuous cryogenic cold-chain, which was kept up until the samples were loaded into the OMNY instrument.^39^

#### Imaging parameters

Ptychographic nanotomography (PXCT) is a coherent diffractive imaging technique, that provides volumetric quantitative electron density of the sample with high spatial resolution.^29^ It relies on illuminating the sample with a coherent, spatially confined x-ray beam. The sample is scanned through the beam ensuring the scanning step size is significantly smaller than the illumination footprint. At each point, an intensity diffraction pattern is collected. Through iterative image reconstruction algorithms,^40^ a projection image of the x-ray sample scattering function is obtained, containing phase-shift and absorption information integrated along the beam path, i.e., the sample thickness. By rotating the sample and repeating the ptychography acquisition, a full tomographic dataset is obtained, from which the electron density distribution within the sample can be retrieved in 3D.^30^ The spatial resolution via ptychography is decoupled from the beam size and step size and is ultimately determined by the largest angle at which diffraction signal can be measured with good signal-to-noise ratio and the accuracy of the sample positioning. PCXT measurements were carried out at the cSAXS beamline (X12SA), Swiss Light Source, Paul Scherrer Institute, Switzerland with the OMNY instrument.^41^ Ptychographic scans were performed at a photon energy of 6.2 keV under cryogenic conditions at 90 K. An Eiger 1.5 M detector, placed 7.2 m downstream of the sample, was used to collect far-field intensity patterns at each scan point. The same ptychography scan was performed at various equiangular orientations, rotating the sample about the vertical axis in the range between 0 and 180°. The exact imaging parameters of each tomogram are listed in Supplementary Table 2.

The ptychographic reconstructions were carried using 300 iterations of the difference-map algorithm,^42^ followed by 300 iterations maximum-likelihood refinement,^42^ resulting in a reconstructed pixel size of 38.5 nm (Supporting Figs. 11).^40^ The projections were further processed and aligned using in-house scripts.^43^ In order to provide a first estimate of the spatial resolution a classical approach was used. The datasets were splitted angularly into two separately reconstructed datasets and compared using the Fourier shell correlation (FSC) approach.^44^ In this way, the half-bit half-period resolution was estimated as ∼53 nm, for a detailed discussion on the resolution of PXCT in the datasets presented see **Supplementary discussion on the spatial resolution of PXCT**. The corresponding FSC curves are presented in Supplementary Fig. 9 and the detailed FSC figures are tabulated in Supplementary Table 2.

Some residual artefacts, likely caused either from misalignment or radiation-induced effects were observed in the tomogram of the cells at stage 4 (Supplementary Movies 4, 5 and 9). Using the glass capillary as a reference, its morphology and electron density were comparable with those of the other tomograms (0.64 – 0.66 n_e_Å^-3^). We therefore considered that these artefacts did not significantly affect the electron density or the morphology of the intracellular structures in the coccoliths in this sample.

#### Segmentation of tomograms

The reconstructed cryoPXCT tomograms gave quantitative electron density values (n_e_Å^-3^).^29^ All segmentation was carried out on Avizo software (ThermoFisher Scientific). For segmentation of electron-dense structures, a minimum electron density threshold value of 0.37 n_e_Å^-3^ was chosen to determine where calcium ions were located. Localized and well-defined regions of electron density above 0.43 n_e_Å^-3^ were often present within the larger compartments with electron density values between 0.37-0.43 n_e_Å^-3^. Those well-defined, denser regions, were segmented separately from the compartment that contained them.

### Nano-scanning X-ray Fluorescence (nXRF) imaging

Measurements were conducted at I-14 X-ray nanoprobe beamline of the Diamond Light Source.^45^ Samples were dried on Si_3_N_4_ membranes and measured at ambient pressure and temperature. An incident photon energy of 8 keV was used with a focused X-ray beam size of ∼60 nm (FWHM). A raster scanning step size of 100□nm was used with a dwell time of 50□ms to collect high-resolution X-ray fluorescence (XRF) maps. XRF from the sample was collected in front of the sample using a four-element silicon drift detector (RaySpec, UK). A photon-counting Merlin detector (Quantum Detectors, UK) was used in transmission configuration to collect scattered X-rays for differential phase contrast (DPC) and phase gradient (PG) images. For semi-quantitative analysis of elemental composition, an AXO XRF standard (AXO DRESDEN GmbH) was measured using identical scanning parameters to coccolithophore samples. The summed XRF spectrum of the AXO standard was used to calibrate the elemental concentrations of P, S, Ca, Mn and Fe for semi-quantitative analysis with reported quantities in fg µm^-2^. PyMCA 5.6.7 software was used to energy calibrate XRF spectra, background subtract spectra, fit XRF sum-spectra, calibrate XRF counts to elemental quantities with AXO XRF standard material and export XRF maps for individual emission lines (e.g., Ca Kα). ImageJ was used to further render XRF maps.

## Acknowledgements

Funding for this research was provided by NERC through an E4 DTP studentship (NE/S007407/1) to F.N. and R.W. and by the European Research Council (European Union’s Horizon H2020 research and innovation program grant agreements No 724881 to V.C. and T.G. Support was provided by the French National Research Agency (microCOCCO: ANR-21-CE42-0022) to D.M.C. CryoPXCT was performed at the coherent small-angle x-ray scattering (cSAXS) beamline at the Swiss Light Source at the Paul Scherrer Institute, Proposals 20181865 and 20200711. We acknowledge Diamond Light Source for time on Beamline I14 under Proposals MG23602 and MG28868. We thank Manfred Burghammer for his help with the acquisition of cryoPXCT data.

## Author Contributions

F.N. and T.G. conceptualized the study. A.T., T.G, F.N and N.O. prepared cryoPXCT samples. F.N., T.G., M.V., J.I., M.H and M.G-S performed ptychographic tomography experiments. M.V. and J.I. reconstructed the tomography data. A.T. conducted tomogram segmentation and A.T., T.G., V.C. and F.N. analysed the data. D.M.C and A.S. prepared nXRF samples and D.M.C conducted the measurements and data analysis. A.T., D.M.C, T.G, V.C. and F.N. wrote the manuscript. F.N. supervised and R.W. co-supervised the study. All co-authors provided input to the manuscript.

