## Supplementary Information for "Dynamic change of calcium-rich compartments during coccolithophore biomineralization"

**This Supplementary Information Includes:**

- Supplementary Figure 1-9
- Supplementary Table 1-2
- Supplementary discussion on the spatial resolution of PXCT
- Supplementary Movies 1-9
- Supplementary References

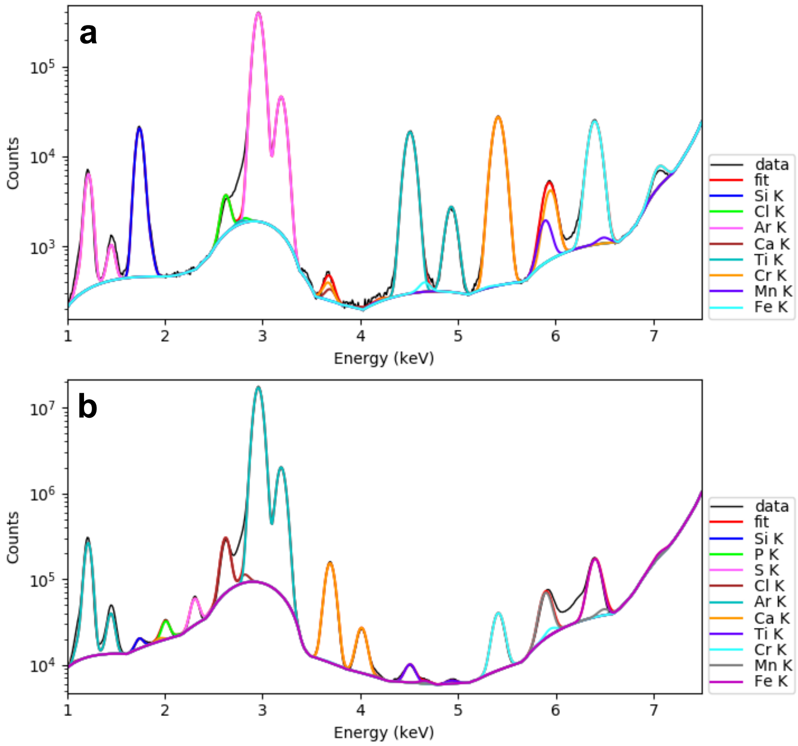


**Supporting Figure 1**. Fitted XRF spectrum of (a) AXO standard material and (b) *C. carterae* harvested after 4 h in Ca-replete medium.

Supplementary Table 1. Semi-quantitative elemental analysis (AXO standard calibrated) of XRF signals for *C. carterae* harvested after 4 h in Ca-replete medium. Values are background subtracted (substrate background) and area-corrected. Average values are calculated from Cell 2 and presented in Figure 2c.

| | Feature | P (fg μm$^{-2}$) | S (fg μm$^{-2}$) | Ca (fg μm$^{-2}$) | Mn (fg μm$^{-2}$) | Fe (fg μm$^{-2}$) |
|---|---|---|---|---|---|---|
| Cell 1 | Compartment | 2.313 | 0.260 | 0.566 | 0.001 | 0.013 |
| | Cell background | 0.184 | 0.782 | 0.147 | 0.002 | 0.017 |
| Cell 2 | Compartment 1 | 9.014 | 1.069 | 2.686 | 0.048 | 0.072 |
| | Compartment 2 | 14.355 | 1.082 | 4.460 | 0.053 | 0.078 |
| | Compartment 3 | 13.418 | 1.443 | 4.131 | 0.053 | 0.079 |
| | Cell background | 1.057 | 1.838 | 0.946 | 0.050 | 0.089 |
| Average | Compartments | 12.3 ± 2.8 | 0.80 ± 0.21 | 3.8 ± 0.9 | 0.051 ± 0.003 | 0.076 ± 0.004 |
| | Cell background | 1.1 ± 0.6 | 1.4 ± 0.2 | 0.9 ± 0.5 | 0.012 ± 0.001 | 0.015 ± 0.001 |

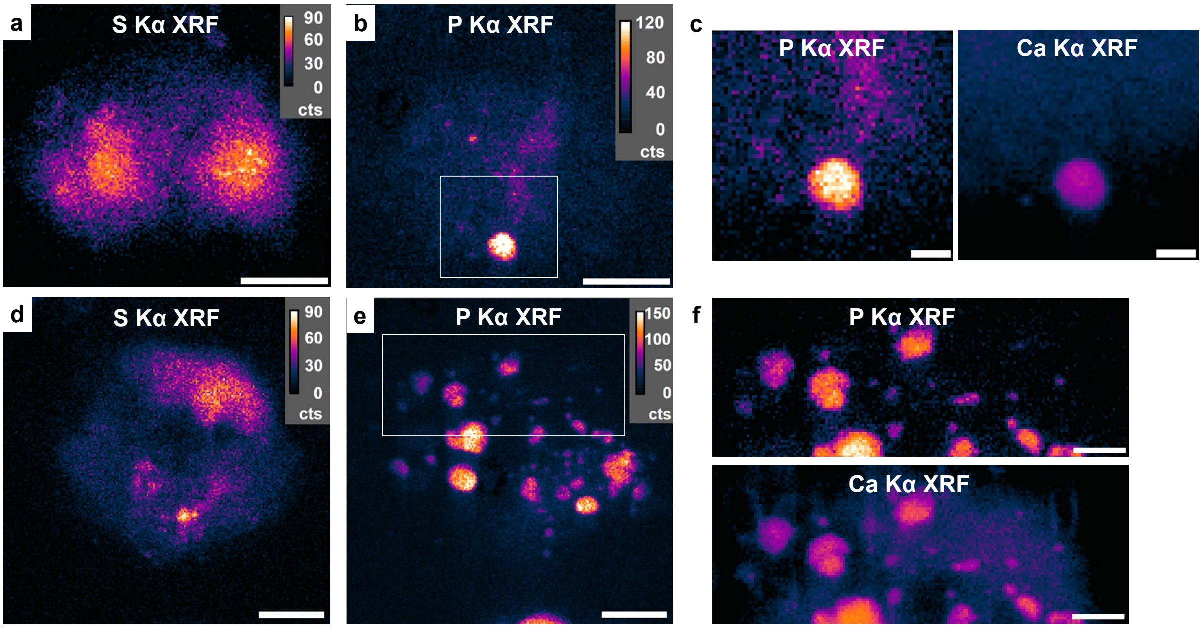


**Supplementary Figure 2**. Nano-XRF maps of *C. carterae* cells (Cell 1 (a-c) and 2 (d-f), as presented in Figure 2) for S Kα XRF and magnified regions (white box in b and e) showing P and Ca Kα XRF signals in log-based intensity (c and f). Co-localization of Ca and P is more evident in log-based intensity. Scale bars: 4 µm (a, b, d, e), 1 µm (c) and 2 µm (f).

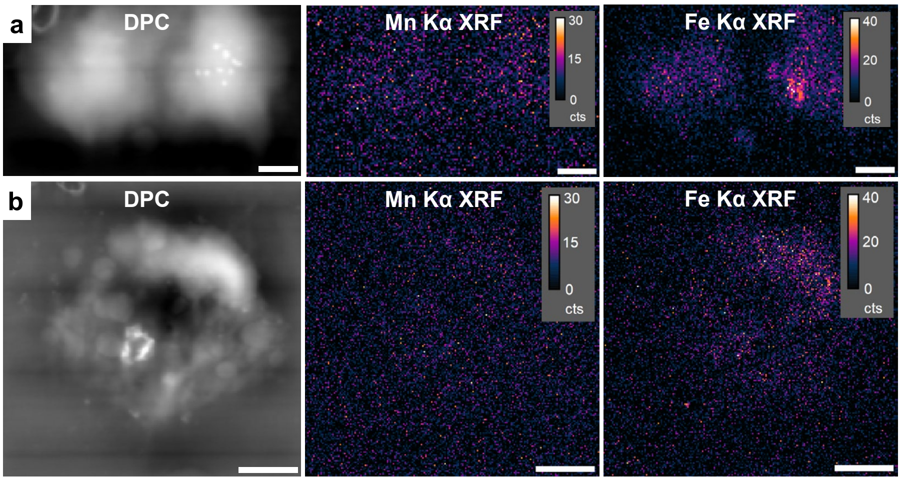


Supplementary Figure 3. Nano-XRF maps of *C. carterae* cells (Cell 1 (a) and 2 (b), as presented in Figure 2) with Mn and Fe Kα XRF signals. These elements show weaker and more disperse signals compared to Ca and P, for example. Cell 1 is cropped to remove metal-rich impurity at the top of the map. Scale bars: 2 µm (a) and 4 µm (b).

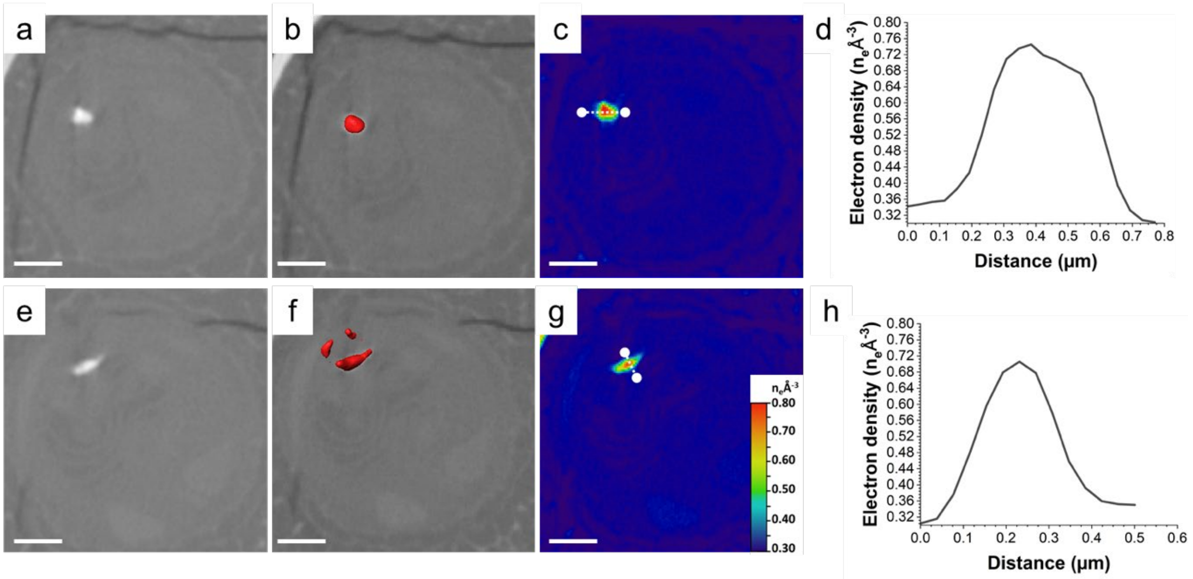


Supplementary **Figure 4: Electron dense calcium-rich structures in a *C. carterae* cell at the 4 h timepoint.** Structures with electron density above 0.70 $n_e$Å$^{-3}$ present in the cell. (a) and (e) show 2D slices through the volume of the cell along the xy plane. (b) and (f) depict the 3D isosurface-rendered reconstructions of the structures overlayed on the 2D slices. The electron density maps of the structures are shown in (c) and (g), together with the electron density profiles (d) and (h), measured along the dotted line in (c) and (g). Scale bars: 2 μm.

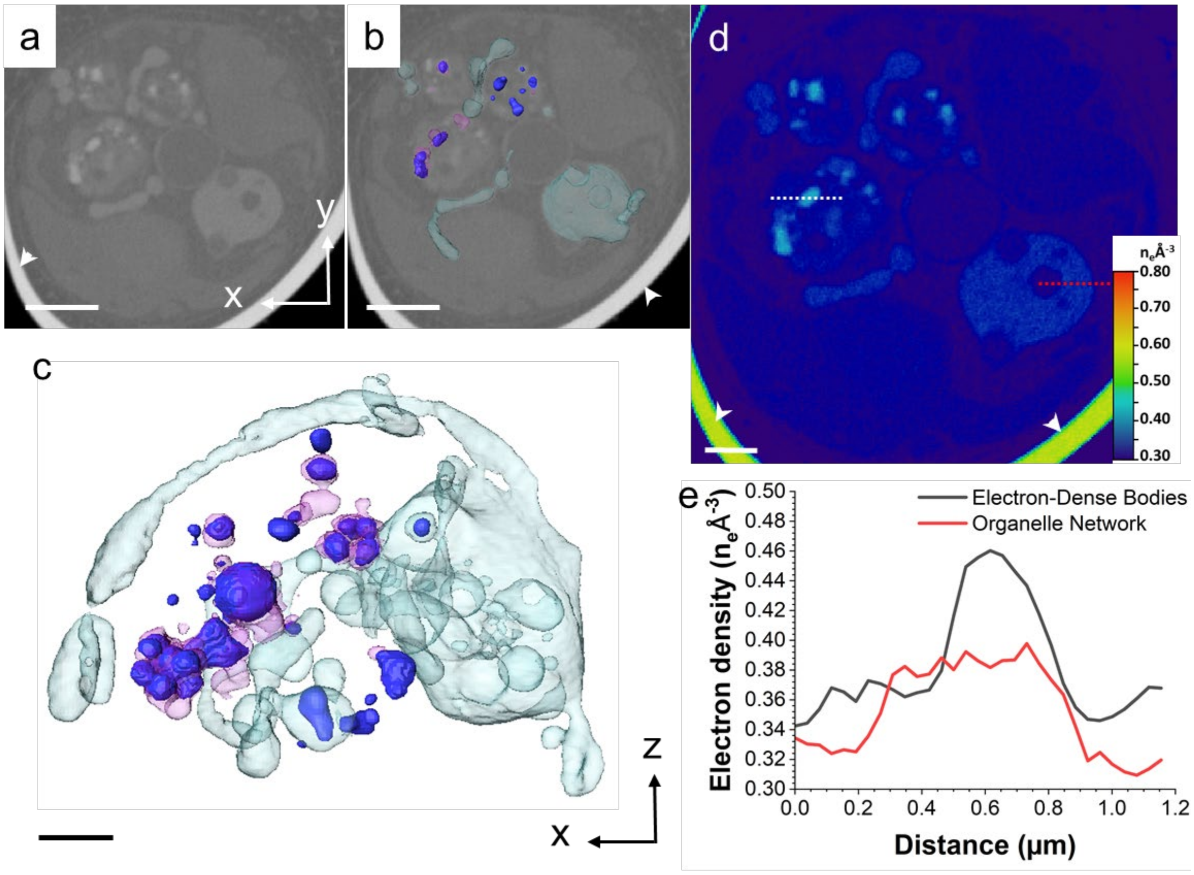


Supplementary **Figure 5: Calcium-rich structures in a non-calcifying *C. carterae* cell at stage 1.** Calcium-rich structures within a cell at stage 1 of calcification were revealed using cryo-PXCT. Even though in calcium-containing medium as the same time as the cell visualized in Figure 3, a cross-section through this cell (a), along with 3D isosurface-rendered reconstructions in (b) and (c), shows electron-dense bodies (dark blue) were still contained within larger compartments (pink), with the organelle network (turquoise) present as well. This is similar to the distribution in the cell present at the 4 h timepoint. (d) Electron density map of a (a). (e) Electron density line profile of the area marked by the white line in d. No intracellular coccoliths were visible in (c), indicating the cell was not calcifying when frozen. Scale bars: 2 µm. Arrowheads indicate the wall of the glass capillary used to store the sample.

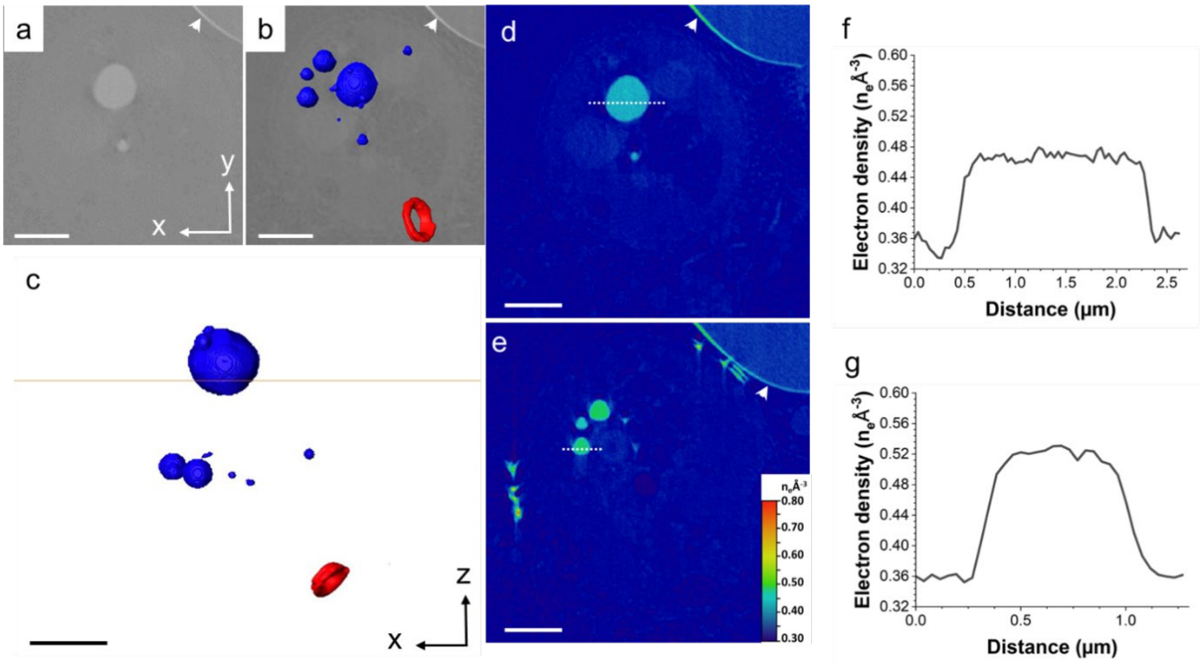


Supplementary **Figure 6: Calcium-rich structures in a *C. carterae* cell at stage 2.** Calcium-rich within another cell left at stage 2 of formation of the coccosphere. Cross-sections (a) and 3D isosurface-rendered reconstructions (b,c) show that globular electron-dense bodies (dark blue – b-c) were present, similar to the cell at the 8H timepoint (Figure 4). Electron density maps of (a) and of a different slice of the volume of the cell are shown in (d) and (e), respectively. (f) and (g) depict the electron density profile measured in the area marked by the dotted line in (d) and (e). Given a related number of external coccoliths in both cells, it is likely they had been producing coccoliths for a similar length of time. Scale bars: 2 µm (a,b; d,e), 1 µm (c). Arrowheads indicate the wall of the glass capillary used to store the sample.

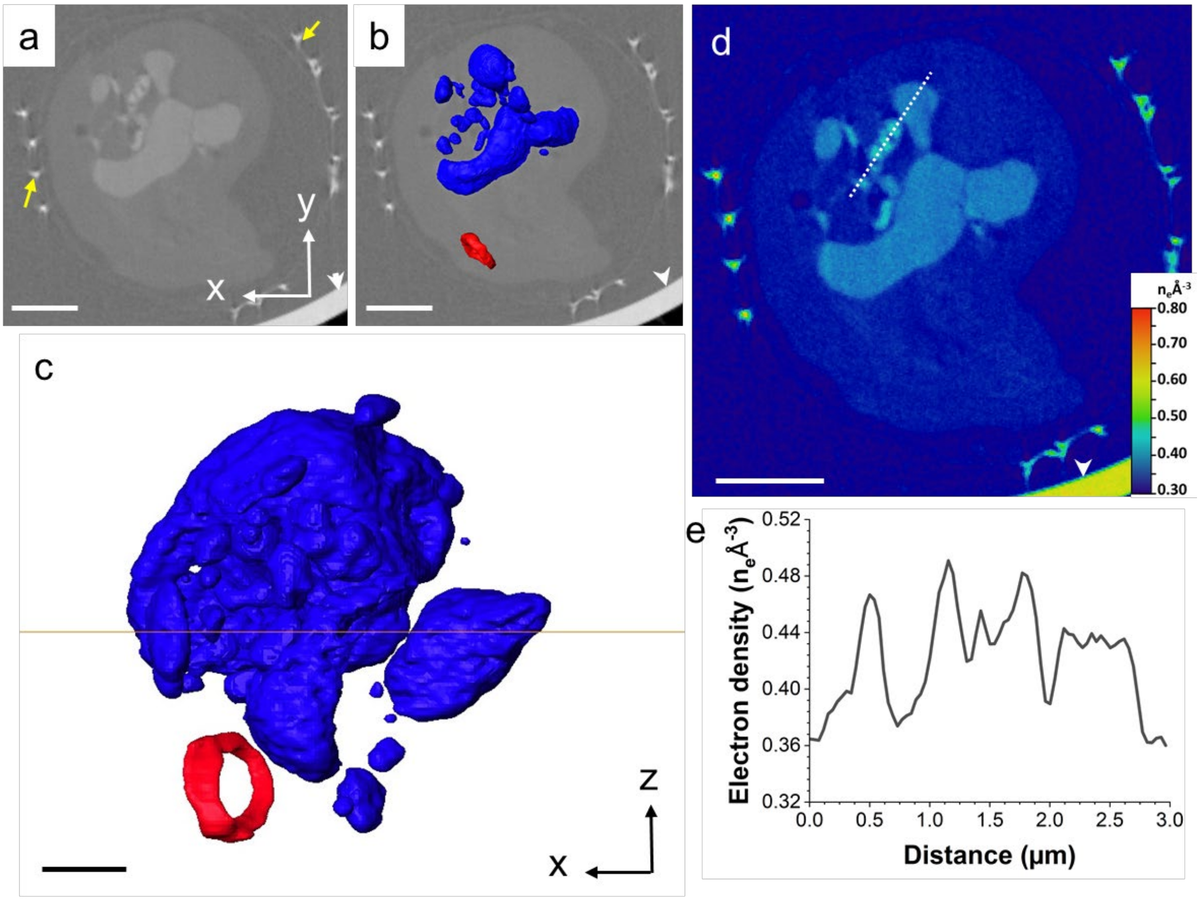


Supplementary **Figure 7: Calcium-rich structures in a *C. carterae* cell at stage 3.** Calcium-rich within a second cell at stage 3 of calcification. (a-e) Cross-sections (a), 3D isosurface-rendered reconstructions (b,c), and electron density map (d) and profile (e), show a similar distribution and calcium content within calcium-containing structures to the cell visualized in Figure 4. Yellow arrows in (a) indicate extracellular mature coccoliths. Scale bars: 2 µm (a,b), 1 µm (c,d). Arrowheads indicate the wall of the glass capillary used to store the sample.

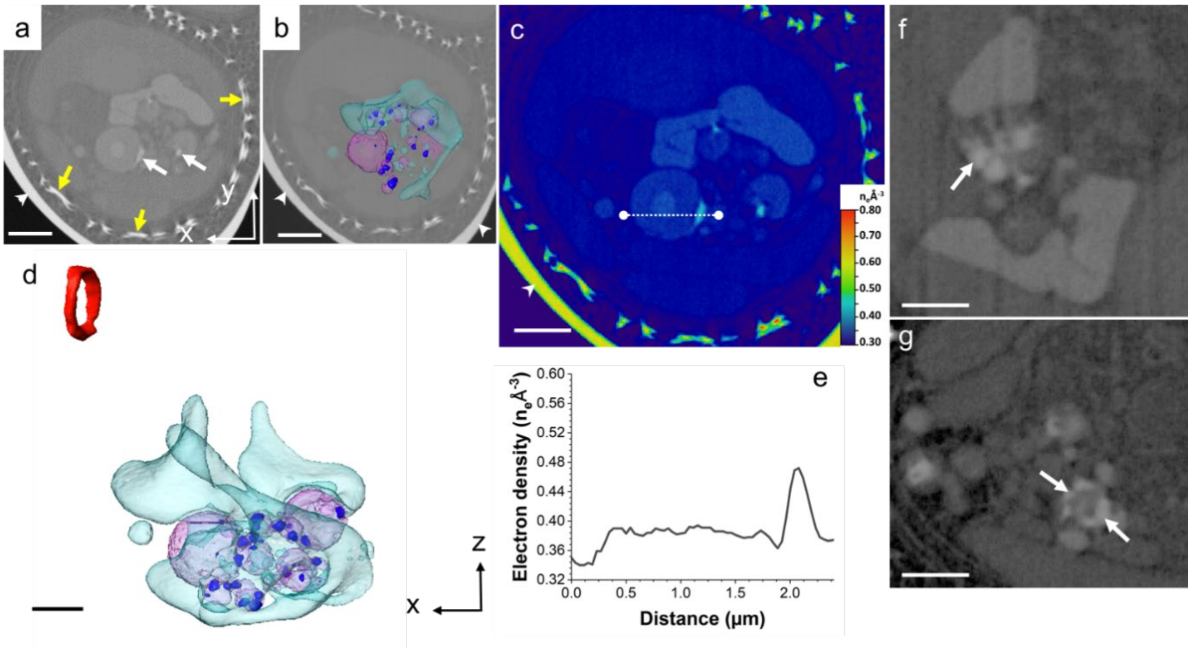


Supplementary **Figure 8: Calcium-rich structures of 3 different *C. carterae* cells at stage 4.** (a). 2D slice through the volume of one cell that, unlike the one represented in Figure 5, is still calcifying. Yellow arrow: Yellow arrows: dense body with electron density in the range of 0.50 $n_e$Å$^{-3}$. The 3D isosurface-rendered visualization of all electron-dense structures is shown overlaid on the 2D slice (b), and in more detail in (c). Visible are: A spherical compartment (pink) with electron density ~0.37 $n_e$Å$^{-3}$ containing bodies with electron density above 0.43 $n_e$Å$^{-3}$ (dark blue); an organelle network with electron density of 0.40 $n_e$Å$^{-3}$ (turquoise); and a forming coccolith (red). (d) Electron density map of a cross-section of the cell depicted in (a-c). (e) Electron density line profiles of the areas marked by the white line in d. (f,g) Two 2D slices from two different cells devoid of forming coccoliths, showing in higher detail electron-dense bodies (white arrows). Scale bars: 2 µm (a-c), 1 µm (d-g). Arrowheads indicate the wall of the glass capillary used to store the sample.

**Supplementary discussion on the spatial resolution of PXCT**

Spatial resolution estimation is a nuanced task. While in conventional (incoherent) microscopy, the resolution can be characterized by the optical transfer function, which describes the resolution properties of the full image, for coherent diffraction-based approaches, other tools have been developed. They are based on the quality of the obtained image.
A first estimate of the spatial resolution is shown hereafter. It is referred to as the Fourier shell correlation (FSC) approach, a widely used method in the ptychography and electron microscopy community. To this end, the dataset were split angularly into two datasets that were separately reconstructed.[46] For FSC analysis, a region inside of the tomogram, without glass capillary walls and surrounding air, was selected. The normalized and angularly averaged cross-correlation coefficients are calculated in the Fourier space, as a function of the spatial frequency. Using the half-bit resolution threshold,[44] a half-period spatial resolution estimate was obtained at 53 nm for the *C. carterae* dataset. The corresponding FSC curves are presented in Supporting Figs. 11 and 12.

While our results from FSC give resolution estimates of about 50 nm, we have however observed that this resolution is supported by edge-sharpness *only* to high-contrast features. Our sample is composed of tiny, weakly scattering elements in the periphery of the coccolithophores, coexisting with strong scattering elements located in the bulk of the coccoliths, these later ones driving the resolution estimation described above. As in any imaging system, low-contrast also results in smaller SNR and therefore can locally present degraded resolution. Examining edge profiles of lower-contrast features, we observed that a density plateau is hardly reached (i.e., at least in one direction of the 3D space, the density maximum is limited to only one pixel). This is illustrated by the electron density line profiles of Fig. 3e (degraded resolution) and Fig. 4e (high resolution). For those elements of the sample, we estimated the spatial resolution to around 200 nm.

In consequence, due to partial-volume effects, the observed electron densities are lower for features that are unresolved or barely resolved. In order to ascertain the results presented in this work are not affected by this issue, we only discuss features for which spatial resolution artefacts can be ruled out. They correspond to the observation of a plateau of the electron density distribution function, i. e., a constant value observed over at least 3x3x3 voxels in 3D. Thereby, the observation of the electron density evolution along the development of the dense-bodies can be unambiguously reported.

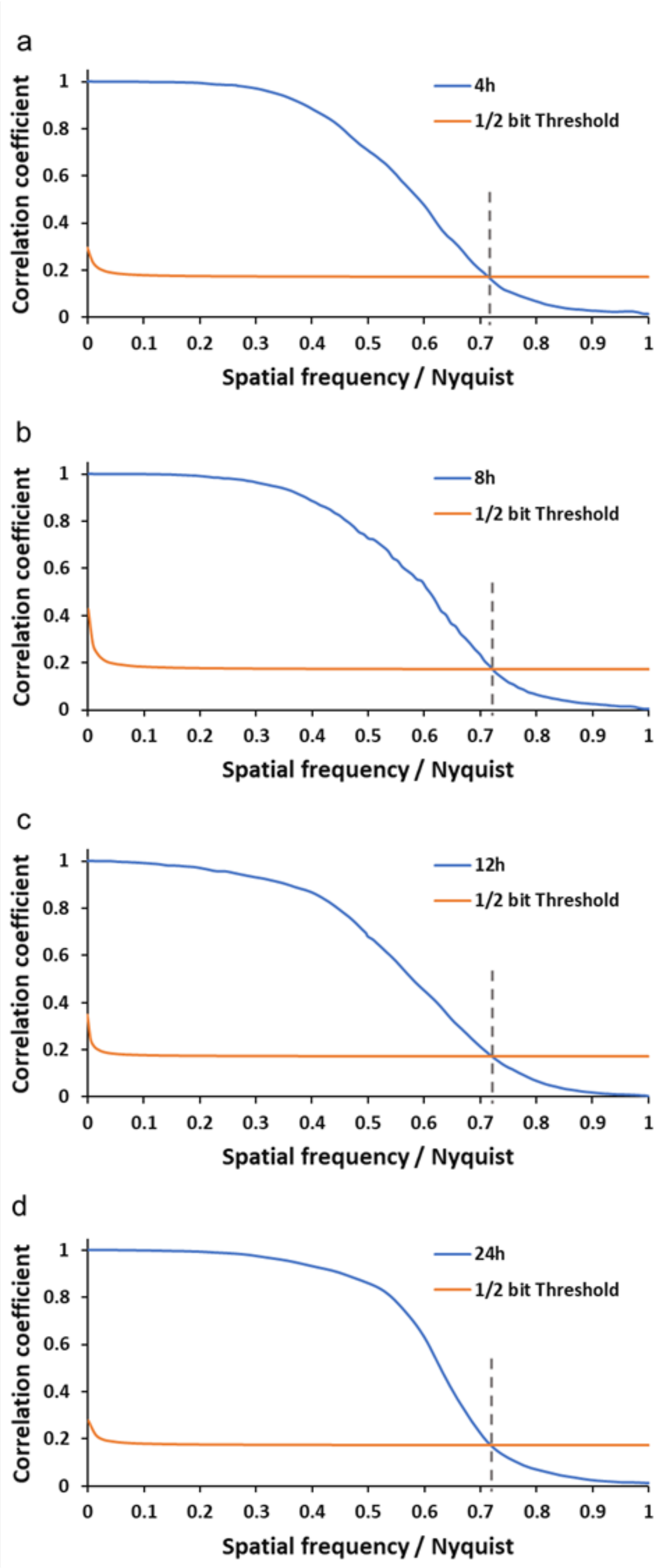


Supplementary **Figure 9**. **Spatial-Resolution of Ptychographic Tomograms obtained from *C. carterae* datasets.** Fourier shell correlation (FSC) line plots of the acquired electron density tomograms. The selected threshold for determining the resolution is the ½ bit criterion. The voxel size for all tomograms is 38.5 nm. The half-period spatial resolution estimate for all samples is ~53 nm. (a) 4 h time point; (b) 8 h time point; (c) 12 h time point; (d) 24 h time point.

Supplementary Table 2. Scan parameters used for the CryoPXCT datasets

| Species | *C. carterae* | | | |
|---|---|---|---|---|
| Time point | 4h | 8h | 12h | 24h |
| Parameter | 286 | 285 | 292 | 293 |
| FOV [µm] | 22 x 17 | 18 x 23 | 38 x 36 | 37 x 25.5 |
| Angles | 465 | 365 | 644 | 800 |
| Step size [µm] | 1.0 | 1.0 | 1.0 | 1.1 |
| Exp time [s] | 0.05 | 0.1 | 0.05 | 0.05 |
| Dose [Gy] | 2.91E+07 | 4.63E+07 | 4.26E+07 | 4.16E+07 |
| FSC Resolution [nm] | 53.9 | 53.4 | 53.7 | 53.4 |

**Supplementary Movie 1:** Orthoslices and volume rendering of a cryoPXCT tomogram of a *C. carterae* cell at stage 1 of the formation of the coccosphere.

**Supplementary Movie 2:** Orthoslices and volume rendering of a cryoPXCT tomogram of a *C. carterae* cell at stage 2 of the formation of the coccosphere.

**Supplementary Movie 3:** Orthoslices and volume rendering of a cryoPXCT tomogram of a *C. carterae* cell at stage 3 of the formation of the coccosphere.

**Supplementary Movies 4 & 9:** Each movie highlights the orthoslices and volume rendering of a cryoPXCT tomogram of one *C. carterae* cell at stage 4 of the formation of the coccosphere, out of 6 that were present in the glass capillary. The tomogram with all 6 cells is shown in **Supplementary Movie 5**.

**Supplementary Movie 6:** Orthoslices and volume rendering of a cryoPXCT tomogram of a second *C. carterae* cell at stage 1 of the formation of the coccosphere.

**Supplementary Movies 7:** Orthoslices and volume rendering of a cryoPXCT tomogram of a second *C. carterae* cell at stage 2 of the formation of the coccosphere.

**Supplementary Movies 8:** Orthoslices and volume rendering of a cryoPXCT tomogram of a second *C. carterae* cell at stage 3 of the formation of the coccosphere.
